## Supplementary figures and images for "Cholinergic and noradrenergic axonal activity contains a behavioral-state signal that is coordinated across the dorsal cortex"

### Figure 1-1

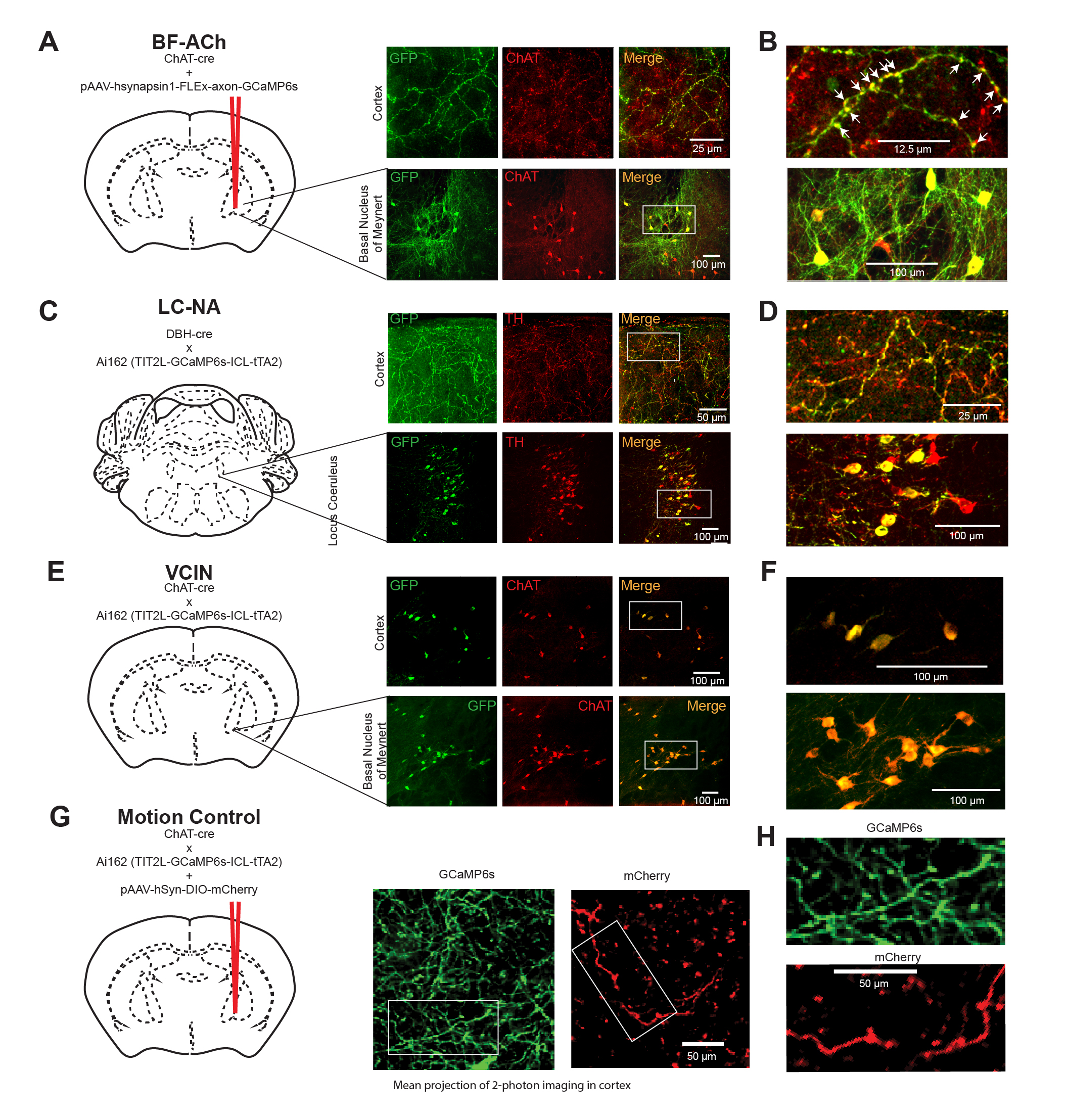

### Figure 2-1

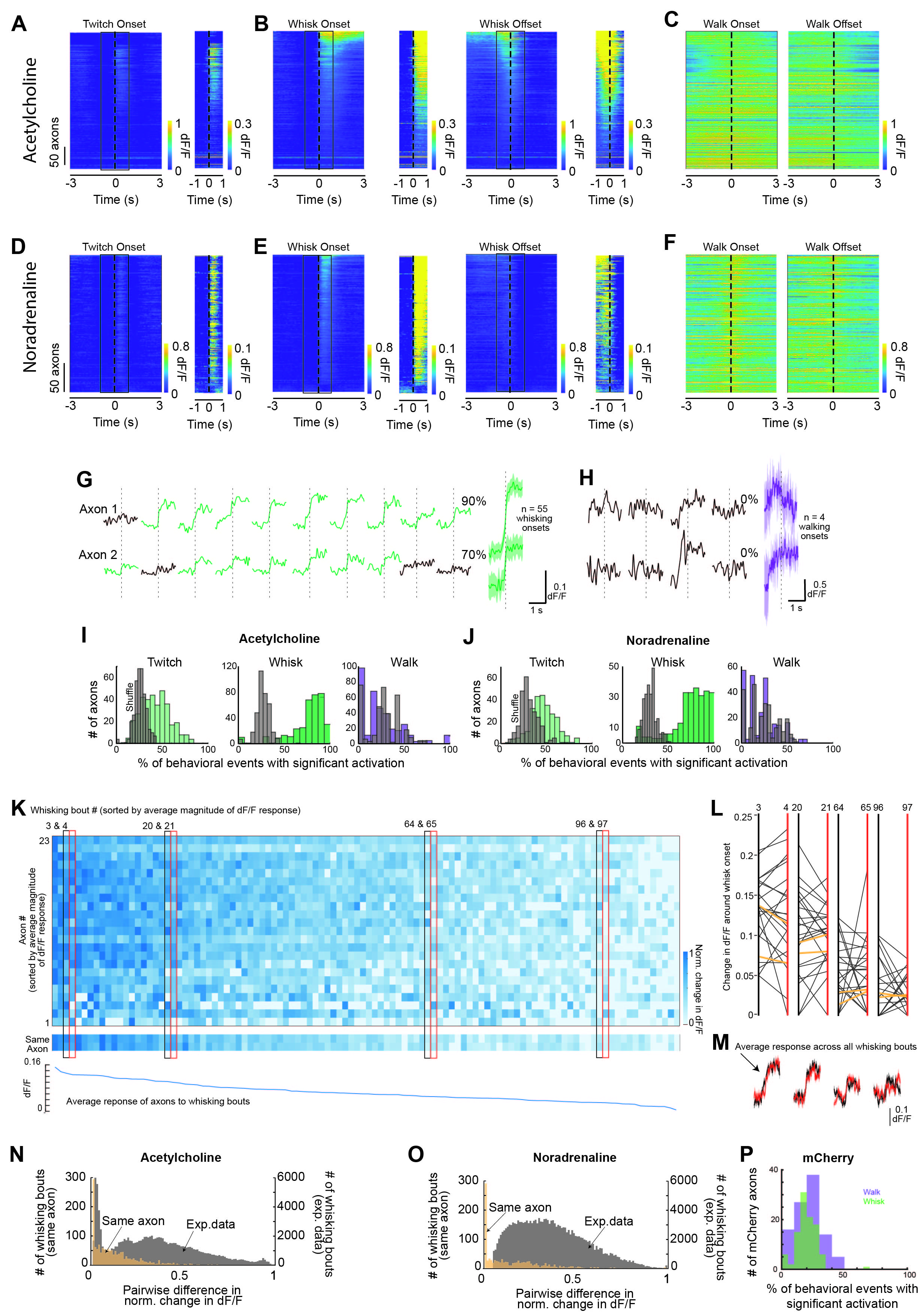

### Figure 2-2

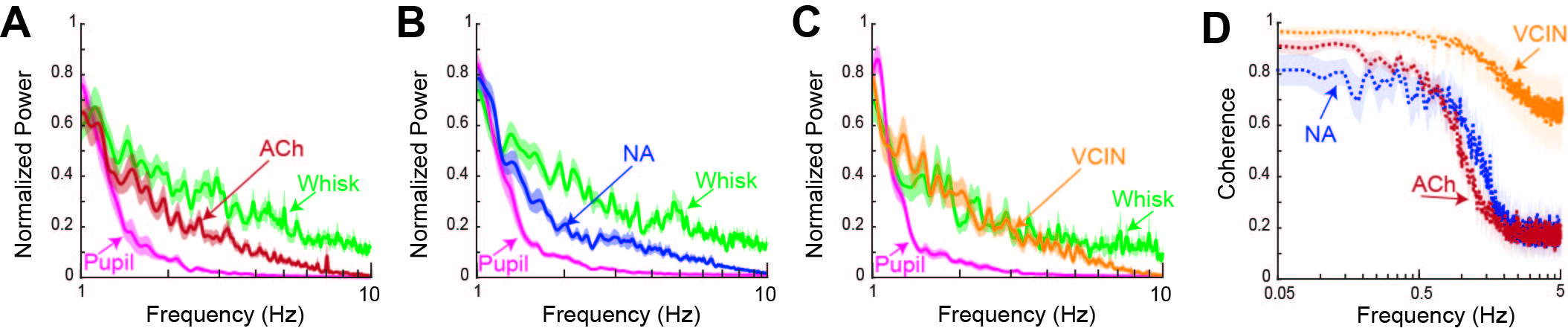

### Figure 2-3

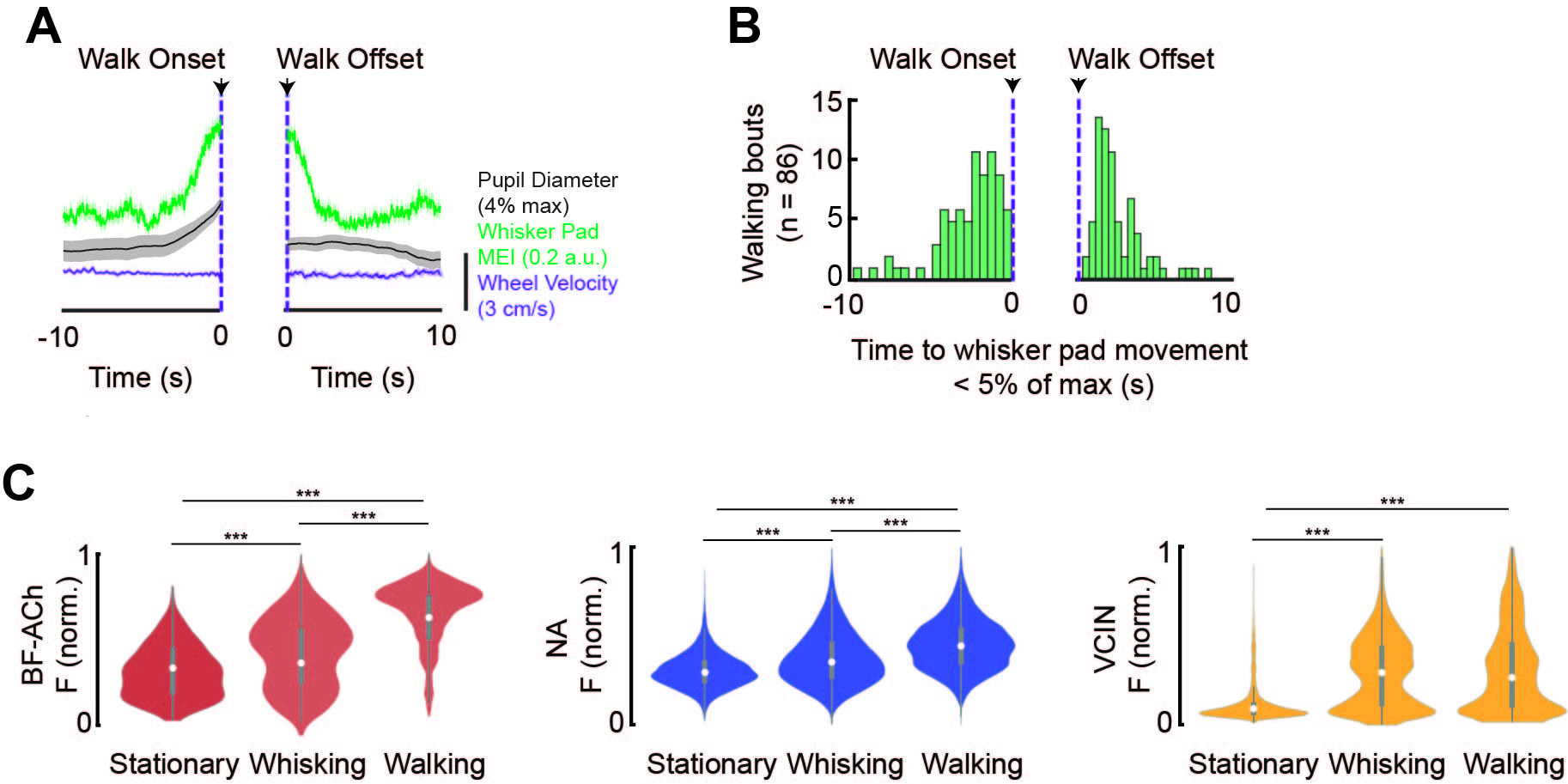

### Figure 3-1

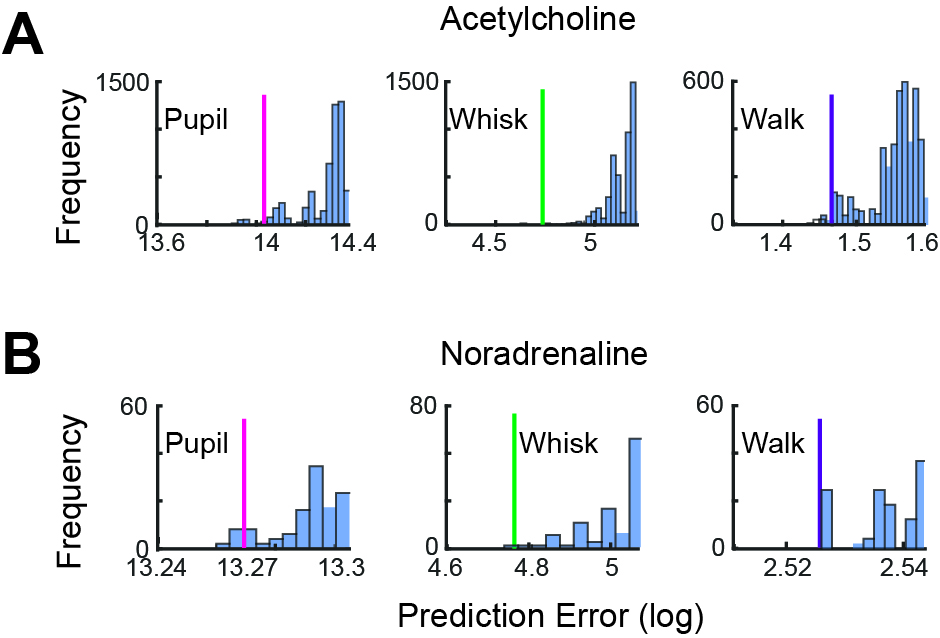

### Figure 4-1

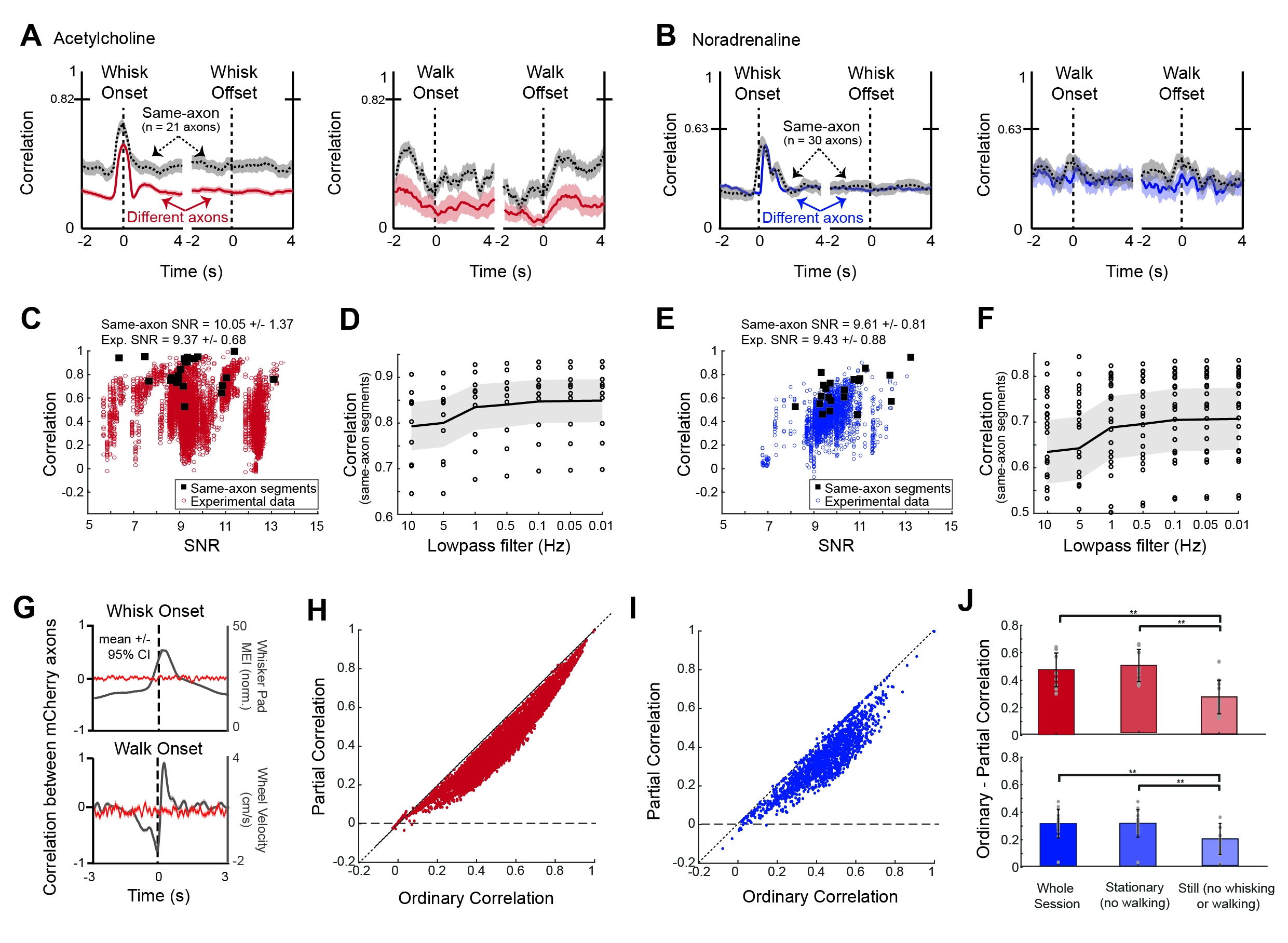
